# ARGONAUTE proteins regulate a specific network of genes through KLF4 in mouse embryonic stem cells

**DOI:** 10.1101/2021.10.18.464771

**Authors:** Madlen Müller, Moritz Schaefer, Tara Fäh, Daniel Spies, Rodrigo Peña-Hernández, Raffaella Santoro, Constance Ciaudo

**Author notes:** These authors contributed equally to this work (Madlen Müller, Moritz Schaefer).

## Abstract

The Argonaute proteins (AGO) are well-known for their essential role in post-transcriptional gene silencing in the microRNA (miRNA) biogenesis pathway. Only two AGOs (AGO1 and AGO2) are expressed in mouse embryonic stem cells (mESCs). The transcriptome of *Ago* mutant mESCs revealed a large and specific set of misregulated genes, compared to other miRNA biogenesis factor mutant cells, suggesting additional functions for the AGOs in stem cells. In this study, we endeavored to understand miRNA-independent roles of the AGOs in gene expression regulation through the integration of multiple datasets. Correlation of *Ago* mutant differential gene expression with ENCODE histone modification data of WT mESCs revealed that affected genes were regulated by the repressive histone modification H3K27me3. We validated this observation by performing chromatin immunoprecipitation followed by sequencing and observed a global loss of H3K27me3 in *Ago* mutant cells. Nevertheless, this reduction explains only a small part of the specific differential gene expression observed in *Ago* mutant mESCs. By integrating chromatin accessibility data in conjunction with prediction of transcription factor binding sites, we identified differential binding for five transcription factors, including KLF4 as a key modulator of more than half of the specific misregulation of gene expression in the absence of AGO proteins. Our findings illustrate that in addition to chromatin state, information about transcription factor binding is more revelatory in understanding the multi-layered mechanism adopted by cells to regulate gene expression. These data also highlight the importance of an integrative approach to unravel the variety of noncanonical functions of AGOs in mESCs.

## INTRODUCTION

The Argonaute (AGO) proteins are well known for their cytoplasmic role in the microRNA (miRNA) pathway, where they are key players involved in miRNA-mediated translational inhibition of target mRNAs (Meister, 2013; Müller et al., 2020). However, more recently, several noncanonical functions, which are not directly linked to the cytoplasmic miRNA-mediated post-transcriptional gene silencing, have been described for the AGO proteins. For instance, several studies have reported nuclear functions for the AGO proteins, such as gene silencing and activation, alternative splicing, chromatin organization and, double-strand break repair (Li et al., 2020; Meister, 2013). Noticeably, most of these functions have been reported in human cancer cell lines and possible noncanonical functions in other systems, such as mouse early development, are only just starting to be understood. In fact, one study reported nuclear localized AGO2 in mouse embryonic stem cells (mESCs) and described a role for AGO2 in post-transcriptional gene silencing within the nucleus (Sarshad et al., 2018). We also recently demonstrated that nuclear AGO1 is linked to the proper distribution of heterochromatin at pericentromeres in mESCs. The depletion of AGO1 disturbed heterochromatin formation at pericentromeric repeats and led to an upregulation of transcripts from those specific regions (Müller et al., 2021).

In mESCs, only two out of the four AGO proteins (AGO1-4) are robustly expressed (Boroviak et al., 2018; Lykke-Andersen et al., 2008; Müller et al., 2020). The deletion of either one of them does not affect the viability of the cells nor their potential to differentiate into the three embryonic germ layers (Ngondo et al., 2018). Interestingly, upon *Ago2* depletion in mESCs, AGO1 levels increase and AGO1 become enriched with miRNAs normally loaded in AGO2, indicating a compensation of AGO2 loss by AGO1 and a redundancy of the two protein functions (Ngondo et al., 2018). However, *Ago2* knockout (KO) mESCs cannot differentiate towards the extraembryonic endoderm, and this defect could not be rescued by overexpressing AGO1 (Ngondo et al., 2018). Thus, even if AGO1 and AGO2 have overlapping functions, especially regarding miRNA-mediated translational inhibition, there seems to be differences in their functional repertoire raising the possibility for additional specialized functions in early embryonic development.

Here, we showed that the comparative analyses of gene expression profiles of mESC lines depleted of key regulators of the miRNA pathway (*Dgcr8*_KO, *Drosha*_KO, *Dicer*_KO and *Ago2&1*_KO) revealed a larger number of specifically differentially expressed genes (DEGs) in *Ago2&1*_KO mESCs that are associated with the positive regulation of RNA metabolic processes. We found that *Ago2&1*_KO mESCs have a global loss of H3K27me3. However, this H3K27me3 reduction in *Ago2&1*_KO mESCs did not significantly impact gene expression. Analyses of chromatin accessibility and chromatin immunoprecipitation followed by sequencing (ChIP-seq) data identified the transcription factors CTCF and KLF4 as major regulators of *Ago2&1*_KO specific DEGs. In summary, our analyses suggested that loss of AGO1 and AGO2 induced changes in H3K27me3 occupancy and gene expression independently of miRNA-pathways due to a misregulation of the key stem cell pluripotency transcription factor, KLF4, revealing a novel axis of gene regulation in mESCs.

## RESULTS

### AGO2&1 regulate the expression of a class of genes in mESCs that do not depend on the miRNA pathway

In order to assess the consequences of the loss of the AGO proteins on mESC gene expression, we integrated available RNA-sequencing (RNA-seq) data from (Schäfer et al., 2021) (Dataset EV1). Previous transcriptomics analyses in multiple *miRNA*_KO (*Dgcr8*_KO, *Drosha*_KO, *Dicer*_KO and *Ago2&1*_KO) and WT mESCs identified many DEGs in all these mutants (Schäfer et al., 2021). Here, we intersected the *Ago2&1*_KO DEGs with DEGs from other *miRNA*_KO mESCs and identified 1793 genes that were specifically misregulated in the AGO1 and AGO2 depleted cells and not altered in any other *miRNA*_KO lines (Fig 1A). Notably, the lack of misregulation in the other mutants ruled out the possibility of direct regulation by miRNAs for these DEGs and there was no overlap with previously reported miRNA targets in mESCs (Fig 1A) (Schäfer et al., 2021). Out of the *Ago2&1*_KO specific DEGs, more than half (1045) were downregulated and 748 were upregulated (log2 fold-change +/− 0.5, *P* value < 0.05, Fig 1B, Dataset EV1). Since a loss of miRNA-mediated repression leads to increased target levels, the high number of downregulated genes again argues against a miRNA-mediated regulation and implies that they might be attributed to AGO-specific functions. Importantly, the deficiency of both AGO1 and AGO2 in mESCs caused this distinct transcriptomic profile, as single *Ago1*_KO and *Ago2*_KO mESCs only had few DEGs, as previously reported (Fig EV1A and B, Dataset EV1) (Müller et al., 2021; Ngondo et al., 2018). Further, the overlap between the single *Ago1*_KO and *Ago2*_KO DEGs was rather small, which supports only a partial compensatory role for AGO1 and AGO2 functions (Fig EV1C) (Müller et al., 2021; Ngondo et al., 2018).

**Figure 1.**
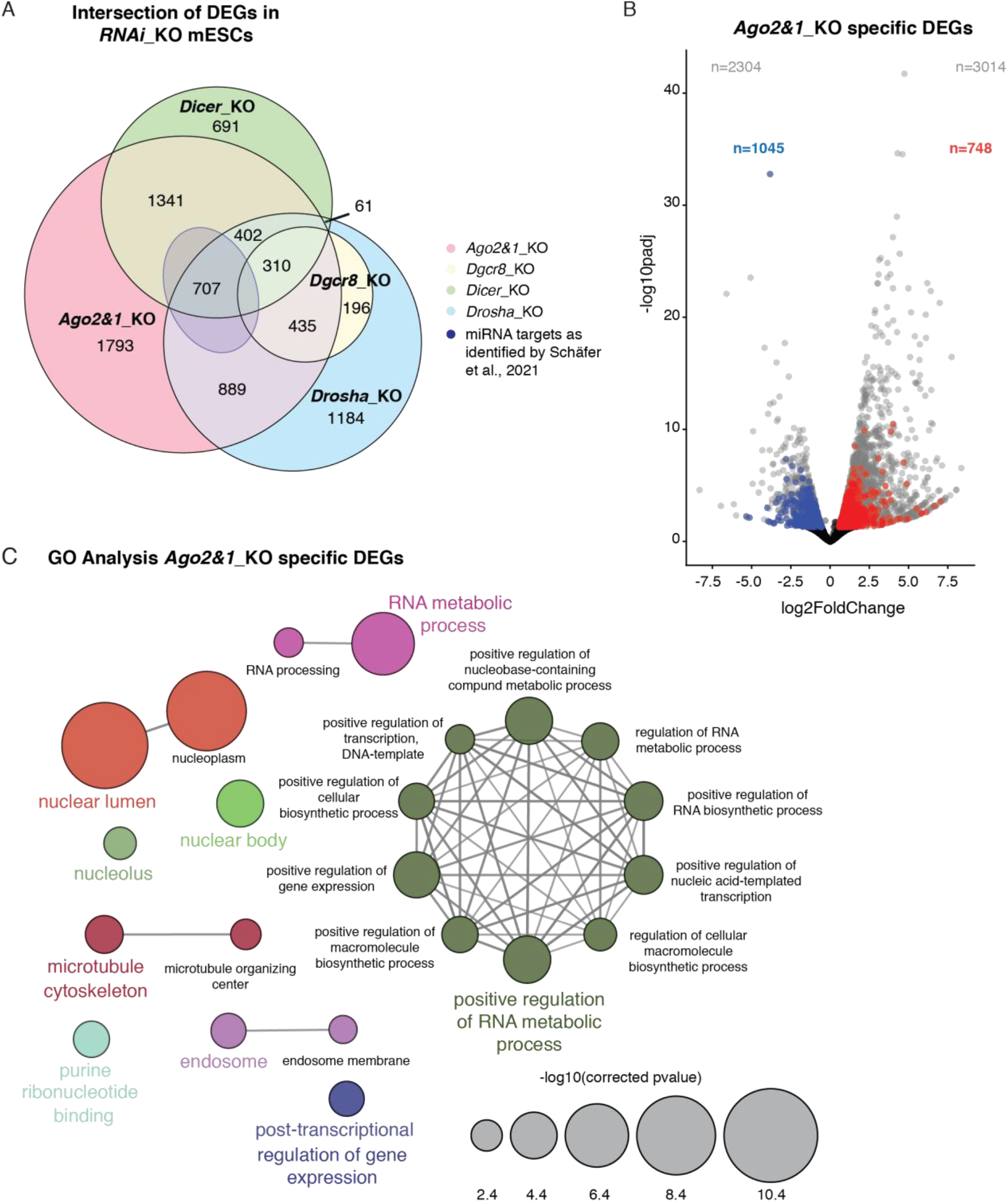
*Ago2&1*_KO mESCs display a distinct transcriptomic profile. (A) Venn diagram representing the overlap of differentially expressed genes (DEGs) from different *miRNA*_KO mESCs (*Dgcr8*_KO, *Drosha*_KO, *Dicer*_KO, *Ago2&1*_KO) and the 707 miRNA targets from (Schäfer et al., 2021). Numbers indicate the gene set sizes of different overlaps. *Ago2&1*_KO mESCs have 1793 specific DEGs (Dataset EV1). (B) Volcano plot showing the *Ago2&1*_KO DEGs and the 1793 specific DEGs. The full set of *Ago2&1*_KO DEGs is shown in gray. Highlighted in red are the *Ago2&1*_KO specific DEGs that are upregulated (748) and in blue the *Ago2&1*_KO specific DEGs that are downregulated (1045). (C) Gene Ontology analysis on the *Ago2&1*_KO the 1793 specific DEGs. The GO analysis has been performed with ClueGO (Bindea et al., 2009). The size of the circles corresponds to their p-value.

In order to understand which pathways are affected in mESCs upon combined loss of AGO1 and AGO2, we performed gene ontology (GO) analysis of *Ago2&1*_KO specific DEGs and found that they were enriched in processes linked to nuclear processes, RNA metabolism, and positive transcriptional gene regulation (Fig 1C). Thus, the function of AGO proteins is not only related to post-transcriptional gene silencing, but also to functions that are probably independent of the miRNA pathway.

### Combined loss of AGO2&1 in mESCs causes global loss of H3K27me3

To determine how the combined depletion of *Ago2&1* affects gene expression in mESCs, we intersected *Ago2&1*_KO RNA-seq and ENCODE histone chromatin immunoprecipitation followed by sequencing (ChIP-seq) from mESCs with the same genetic background (Fig 2A). Specifically, we analyzed the occupancy of histones containing repressive marks (H3K9me3 & H3K27me3), enhancer marks (H3K4me1 & H3K27ac) and active marks (H3K9ac, H3K36me3 & H3K4me3) at *Ago2&1*_KO specifically up- and downregulated genes. As control, we measured the occupancy of modified histones at predicted functional miRNA targets (Schäfer et al., 2021) and expressed genes in mESCs. We detected only minor differences in the occupancy of active histone marks between the *Ago2&1*_KO specifically up- and downregulated genes and the expressed genes in mESCs (Fig 2A). In contrast, we found an enrichment of H3K4me1, H3K27ac, and H3K27me3 at *Ago2&1*_KO specifically upregulated genes compared to downregulated genes and expressed genes in mESCs. Further, the differential H3K27me3 occupancy at *Ago2&1*_KO specifically up- and downregulated genes seemed to be rather correlating with the loss of AGO proteins and not due to a miRNA-mediated regulation, as miRNA target genes (Schäfer et al., 2021) are mainly more enriched in active histone marks than H3K27me3 (Fig 2A). Noticeably, recent results have linked AGO2 with H3K27me3 by showing a reduction of H3K27me3 at certain target loci upon *Ago2* depletion in mESCs (Kelly et al., 2019). To determine whether the combined loss of AGO2&1 affects H3K27me3 levels and gene expression, we first measured and compared the global levels of several histone marks in *Ago2&1*_KO mESCs and WT mESCs by Western Blotting (WB) (Fig 2B). We observed a drastic reduction of H3K27me3 signal in *Ago2&1*_KO mESCs compared to WT mESCs whereas the levels of other modified histones such as H3K9me3, H3K27ac, and H3K4me3 were not particularly affected. These results indicated that the combined loss of AGO2&1 globally decreased H3K27me3 levels. PRC2 is the complex involved in the deposition of H3K27me3 mark (Margueron and Reinberg, 2011). Interestingly, it was recently reported in cancer cells that miRNAs reinforce the repression of Polycomb Repressive Complex 2 (PRC2) transcriptional targets through independent and feed-forward regulatory networks (Shivram et al., 2019). In addition, several members of the PRC2 complex have been shown to be directly regulated by miRNAs in Drosophila (Kennerdell et al., 2018), but not in mESCs deleted for *Dicer*, excluding a direct regulation by miRNAs (Graham et al., 2016). In order to assess the integrity of the PRC2 complex in *Ago2&1*_KO mESCs and the potential contribution of miRNA regulation, we measured the expression of SUZ12 and EZH2, two core proteins of PRC2, in our series of *miRNA*_KO mESC lines. We observed that especially SUZ12 and, to a lesser extent, also EZH2 were specifically downregulated at the protein level (Fig EV2A), but not at the RNA level (Fig EV2B), in *Ago2&1*_KO mESCs. In contrast, and consistent with previous results (Graham et al., 2016), we did not observe changes in SUZ12 and EZH2 at protein and mRNA level in the other *miRNA*_KO mESC lines. These results indicated that AGO1 and AGO2 globally regulate H3K27me3 levels by affecting the protein levels of two key components of PRC2 complex using mechanisms that are independent of miRNA-mediated post-transcriptional pathway.

**Figure 2.**
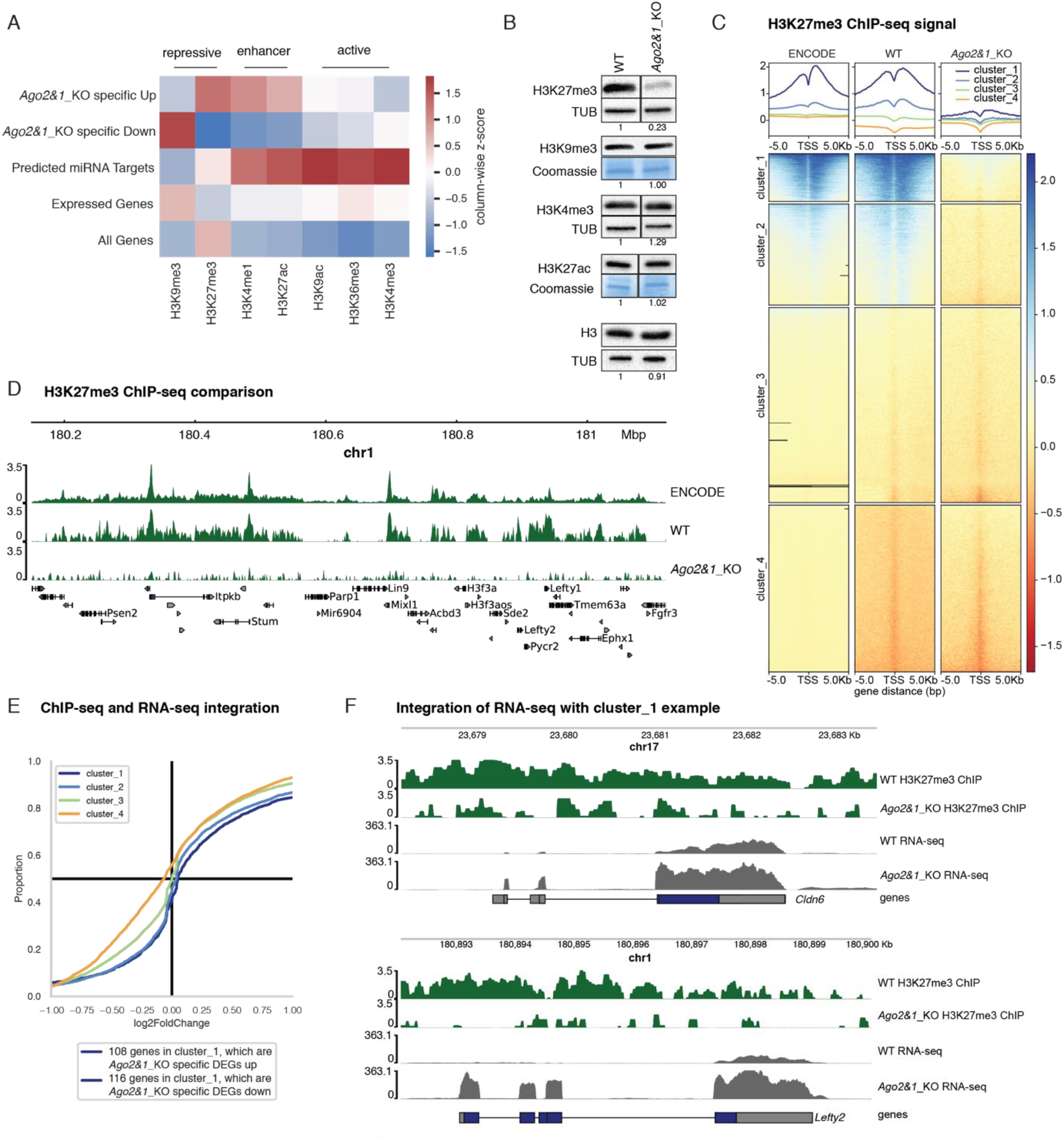
Integration of the *Ago2&1*_KO transcriptome with histone modification datasets. (A) Heatmap showing average histone modification signals at gene regions as derived from ENCODE datasets (see Materials & Methods) for five different gene groups; *Ago2&1*_KO specific up- and downregulated DEGs, 707 predicted functional miRNA target genes from (Schäfer et al., 2021), all expressed genes in mESCs and all annotated genes in mESCs. Histone marks were annotated with their previously described predominant functions (Bannister and Kouzarides, 2011). Repressive histone marks: H3K27me3 and H3K9me3, enhancer marks: H3K4me1 and H3K27ac, activating marks: H3K36me3 and H3K4me3. Columns were individually z-score normalized. (B) Representative Western blots for H3K27me3, H3K9me3, H3K4me3, H3K27ac and H3 in WT and *Ago2&1*_KO mESCs out of n=3 independent experiments. Tubulin (TUB) and Coomassie were used as a loading control. Quantification for each individual experiment is shown below the blot. (C) H3K27me3 ChIP-seq heatmaps of the transcription start sites (TSS) for the full set of annotated mouse transcripts. Shown is one replicate per condition/experiment for ENCODE WT (Davis et al., 2018; Dunham et al., 2012), WT and *Ago2&1*_KO samples (see Materials & Methods). The shown transcript regions have been divided into four different clusters using k-means clustering (Dataset EV2). Shown are +/−5kb from the transcriptional start site (TSS). (D) Genome browser view of a region derived from transcripts from cluster_1. ChIP-seq coverage signals are shown for the three samples from (C) along with annotated genes in that region (bottom). (E) Cumulative Distribution Function (CDF) plot showing the differential expression in *Ago2&1*_KO versus WT of genes associated with the different ChIP peak clusters identified in the ENCODE and WT datasets. The x-axis represents the log2FoldChange of the *Ago2&1*_KO versus WT RNA-seq and the y-axis the cumulative proportion over the full set of log2FoldChanges for each cluster. (F) Genome browser view of two example genes from cluster_1 (Cldn6 and Lefty2) showing both ChIP-seq and RNA-seq coverages for WT and *Ago2&1*_KO (one sample per experiment and condition). The y-axis represents the deposition of H3K27me3 (for the ChIP-seq in green) or the expression levels of the RNA-seq in gray.

Due to the differential H3K27me3 occupancy at *Ago2&1*_KO specifically up-versus downregulated genes (Fig 2A) and the drastic global reduction of H3K27me3 in *Ago2&1*_KO mESCs (Fig 2B) we analyze whether differential H3K27me3 levels affect gene expression. First, we performed H3K27me3 ChIP-seq in WT and *Ago2&1*_KO mESCs (Fig 2C). To validate our H3K27me3 ChIP-seq in WT mESCs, we compared it to available H3K27me3 ChIP-seq data from ENCODE (Davis et al., 2018; Dunham et al., 2012), observing a clear overlap between the two ChIP-seq data (Fig 2C and D). Consistent with the WB analysis, the ChIP-seq data confirmed a genome-wide loss of H3K27me3 in *Ago2&1*_KO mESCs. In order to determine whether H3K27me3 loss correlates with changes in the expression of the *Ago2&1*_KO DEGs, we clustered ChIP-seq levels at transcript regions using k-means (k=4) clustering (Dataset EV2) (Ramírez et al., 2016). The first cluster (cluster_1) represents the genes most strongly enriched in H3K27me3 in WT and highly overlaps with known bivalent genes in mESCs (Asenjo et al., 2020) (Fig 2C, EV2C, Dataset EV2), while the second, third and fourth clusters represent transcripts with minor H3K27me3 levels.

We expected the loss of the repressive histone mark H3K27me3 to lead to an observable upregulation of associated genes. Indeed, genes from cluster_1 and cluster_2, which show the strongest H3K27me3 levels and therefore the strongest loss thereof, show a tendency to be upregulated (Fig 2E and F). In contrast, genes from cluster_3 and cluster_4, which are not marked with H3K27me3 in WT, show no enrichment for upregulation (Fig. 2E, EV2F). However, we found that only 14% of the upregulated genes and 11% of the downregulated genes belong to cluster_1 (Fig 2E, EV2D and EV2E, Dataset EV5). These results suggest that the loss of H3K27me3 has a minor impact on the expression of *Ago2&1*_KO specific DEGs. Thus, other pathways might be specifically affected in the absence of the AGO proteins to explain the specific DEGs.

### Loss of AGO2&1 affects chromatin accessibility in mESCs

To determine how AGO2&1 affect gene expression, we measured chromatin accessibility by ATAC-seq (Buenrostro et al., 2013). We identified 3137 regions exhibiting significant differential accessibility (DA) in *Ago2&1*_KO versus WT (2290 sites with increased accessibility, 847 regions with decreased accessibility) (Fig 3A, EV3A, Dataset EV3). In contrast, only minor differences in chromatin accessibility were observed in single AGO mutants, suggesting that only the combined loss of AGO2&1 can affect chromatin structure of mESCs (Fig EV3B and C). These observations are in parallel with the changes observed at the transcriptomic level in these mutant cell lines, where *Ago2&1*_KO, but not the single KO mutants, exhibited a strongly perturbed transcriptome (Fig EV1A and B). This suggests that the *Ago2&1*_KO specific DEGs might at least be partially explained by changes in chromatin accessibility.

**Figure 3.**
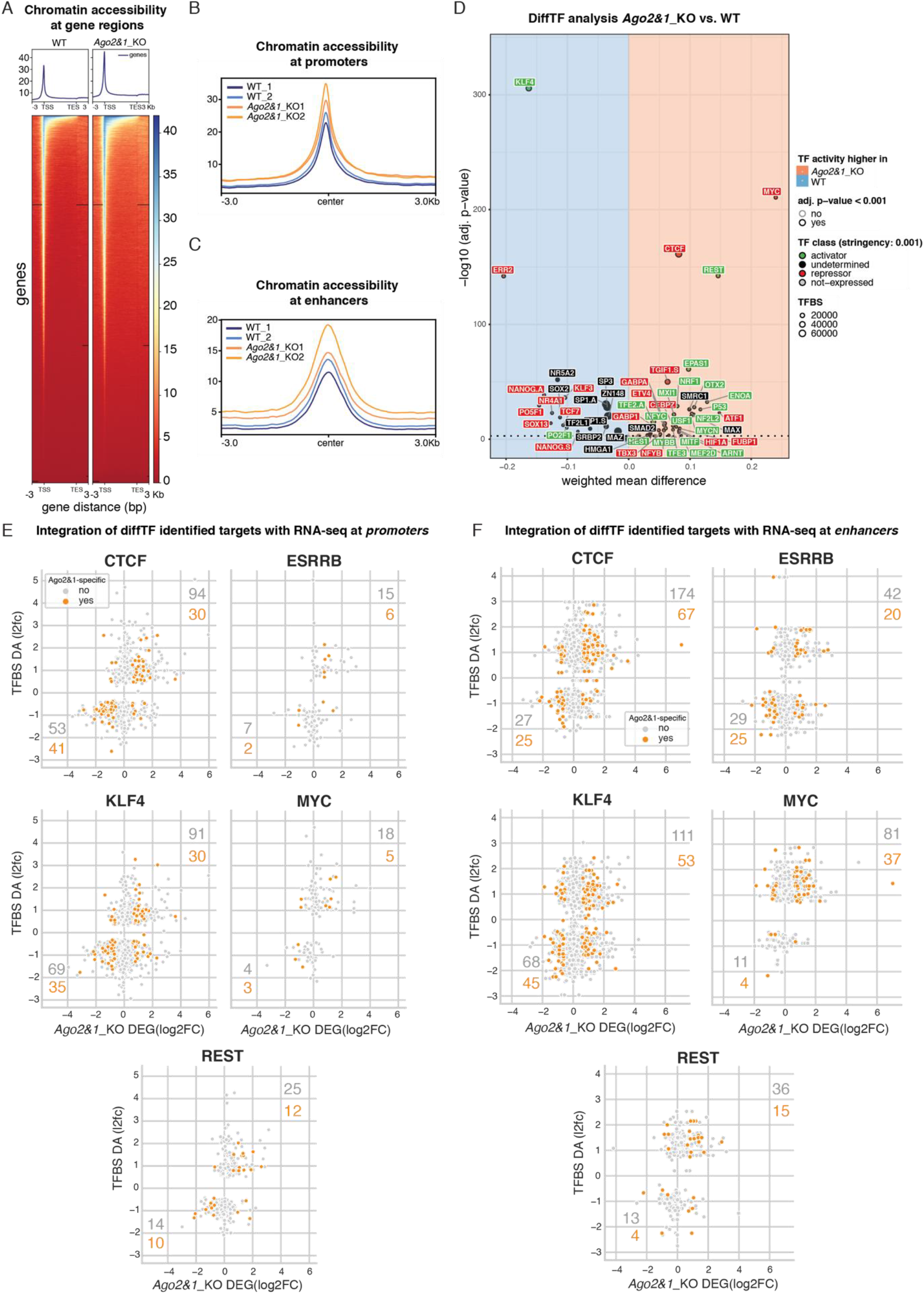
Integration of the *Ago2&1*_KO transcriptome with chromatin accessibility and TF binding. (A) Heatmap and profile plots for WT and *Ago2&1*_KO chromatin accessibility as assessed by ATAC-seq. Transcription start sites (TSS) to transcription end sites (TES) with 3kbp margins are shown for the full set of annotated transcripts. Representative samples of biological duplicates are shown. (B, C) Average signal of chromatin accessibility at TSS/promoter regions (B) and enhancer regions (C), as annotated by (González-Ramírez et al., 2021) for WT and *Ago2&1*_KO samples as assessed by ATAC-seq. (D) Volcano plot of differential chromatin accessibility (DA) at transcription factor (TF) binding sites (BS) for 88 expressed TFs as computed by diffTF (Berest et al., 2019) (Dataset EV4). x-axis shows the difference in chromatin accessibility between *Ago2&1*_KO and WT ATAC-seq samples, where the red area denotes an increase in chromatin accessibility in the *Ago2&1*_KO samples and the blue area a decrease. TFs are annotated as activators (green) or repressors (red) according to the DA at their binding sites and their expression levels, based on RNA-seq data. (E, F) Scatterplots showing differential expression (RNA-seq) versus DA of potential target genes associated with TFBS from (D) for the five TFs with most significant DA binding sites from (D). TFBS were associated with genes by promoter proximity (E, <1kbp distance to TSS) or enhancer proximity (F) (González-Ramírez et al., 2021). Genes are denoted in orange if they are *Ago2&1*_KO specific DEGs.

Next, we retrieved genes that were associated with statistically significant DA regions (Dataset EV3) at their promoter regions (transcription start site (TSS)-distance < 1kbp) and studied their differential expression (DE as indicated by log2 fold-change) in *Ago2&1*_KO mESCs. We observed that the DE of genes with increased chromatin accessibility showed a strong enrichment for increased expression, while genes with decreased chromatin accessibility showed a tendency for decreased expression levels (Fig. EV3D). The difference between the DE distributions of the two groups showed statistical significance (t-test p<1.8e-5). Nevertheless, out of the 384 genes that showed increased chromatin accessibility, only 21 were *Ago2&1*_KO specific DEGs and only 14 of them were upregulated (Fig EV3D and E, Dataset EV3 and EV5). Further, none of *Ago2&1*_KO specific downregulated genes showed significant decrease in promoter accessibility (Fig EV3E and G). Thus, chromatin opening at gene promoter regions alone is not sufficient to explain the specific differential gene expression observed in *Ago2&1*_KO mESCs.

### Transcription factors explain an important part of the specific gene expression in *Ago2&1*_KO mESCs

While chromatin accessibility can influence transcription factor (TF) binding (Spitz and Furlong, 2012), TF binding has conversely been suggested to modulate chromatin accessibility in many cases (Baek et al., 2017), potentially affecting the expression of associated genes. To assess whether TFs might mediate differential chromatin accessibility, we analyzed the DA specifically at promoter (TSS-distance < 1kbp) and enhancer regions (González-Ramírez et al., 2021). We observed an increased chromatin accessibility at these regulatory elements in *Ago2&1*_KO compared to WT mESCs (Fig 3B and C), suggesting increased TF activity. Next, we integrated our chromatin accessibility data with motif-based TF binding site (TFBS) predictions using diffTF (Fig 3D, Dataset EV4) (Berest et al., 2019). Notably, five TFs (CTCF, KLF4, ERR2, REST and MYC) showed highly significant differential binding (Fig 3D, Dataset EV4), which might impact gene expression of their downstream targets in *Ago2&1*_KO mESCs. To assess this impact, we further associated the motif-based TFBS predictions from diffTF with genes, based on promoter-(TSS-distance < 1kbp) and enhancer-proximity (González-Ramírez et al., 2021), and compared DA at TFBS with differential gene expression (Fig 3E and F, Datasets EV4 and EV5). For most TFs, a notable positive correlation between DA and DE was observable, indicating that differential binding of TFs indeed seemed to affect gene expression, thus potentially explaining *Ago2&1*_KO specific DEGs. Combined, the five identified TFs positively correlate with 289 *Ago2&1*_KO specific DEGs (152 up- and 137 downregulated), from which CTCF and KLF4 contributed the most (149 and 147 genes respectively, Fig 3E and F, Dataset EV4 and EV5).

In summary, we observed an increase in chromatin accessibility at promoters and enhancers in *Ago2&1*_KO mESCs and identified five TFs showing highly significant DA at their predicted binding sites. Association of DA at TFBS with proximal genes correlated with around 17% of the *Ago2&1*_KO specific DEGs of which KLF4 and CTCF binding sites correspond to the largest portion (>13%).

### KLF4 regulates the majority of *Ago2&1*_KO specific DEGs

Motivated by the relatively large number of genes predicted as KLF4 and CTCF targets, we further investigated these two TFs. CTCF is well-known for its role in chromatin looping and organization (Oudelaar and Higgs, 2021). More recently, KLF4 has also been linked to chromatin organization during the reprogramming of mouse embryonic fibroblasts (MEFs) to induced pluripotent stem cells (iPSCs) by conferring enhancer-promoter contacts (Di Giammartino et al., 2019). Thus, altering CTCF and KLF4 levels or their binding site accessibility might affect the interaction of regulatory elements and the underlying gene expression. We did not observe a change in *Ctcf* expression at RNA nor protein levels by qPCR and by WB in *Ago2&1*_KO mESCs (Fig EV4A and B). In contrast, KLF4 was strongly downregulated in *Ago2&1*_KO compared to WT mESCs (Fig 4A, Dataset EV1), suggesting that the decrease in KLF4 levels might affect gene expression in *Ago2&1*_KO mESCs. Despite the strong differences in the observed misregulation between the two TFs, they had been associated with a similar number of target genes by our TFBS motif and ATAC-seq-based analysis. ChIP-seq has been recently performed in mESCs for both TFs allowing us to assess and study their targets with higher accuracy (Di Giammartino et al., 2019; Nora et al., 2017).

**Figure 4.**
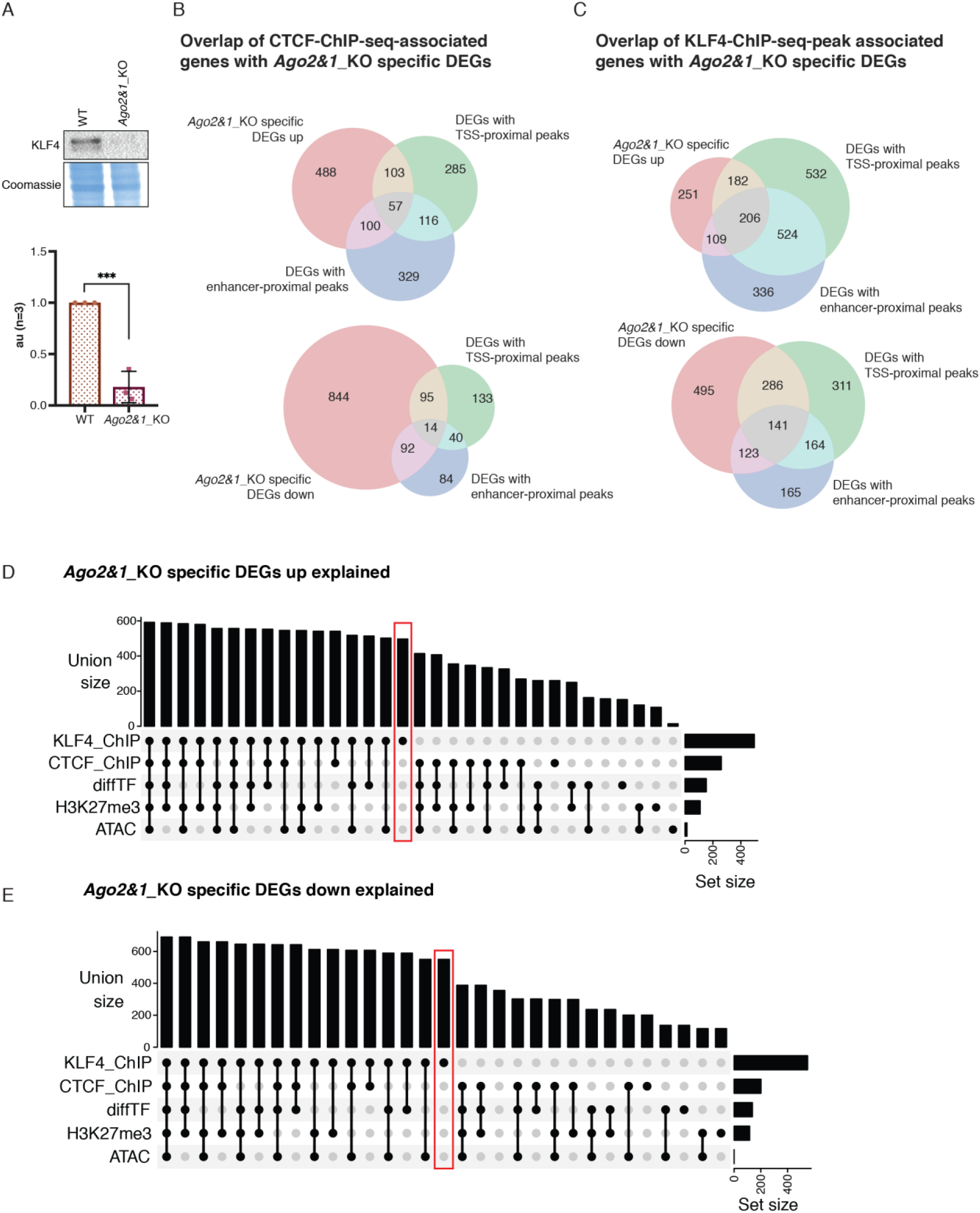
Identification of KLF4-/CTCF-targets and complete integration. (A) Representative Western blot (top) for KLF4 in WT and *Ago2&1*_KO mESCs and quantification (bottom) out of n=3 independent experiments, ***p<0.001, unpaired t-test. (B, C) Venn diagrams of genes identified by CTCF (B) and KLF4 (C) ChIP-seq peak analysis with *Ago2&1*_KO specific DEGs. ChIP-seq peaks were associated with genes by promoter proximity (transcription start site (TSS) < 1000bp, green circle) and by enhancer-proximity (overlap with annotated enhancers by (González-Ramírez et al., 2021)). Different Venn diagrams are shown for upregulated (top) and downregulated (bottom) gene sets and only statistically significant *Ago2&1*_KO differentially expressed genes (DEGs) are considered. (D, E) UpSet plots showing the number *Ago2&1*_KO specific upregulated (D) and downregulated (E) genes explained by one or multiple combined regulatory mechanisms as studied in this paper (Dataset EV5). The dot-connected lines indicate which gene explanation sets were combined and the bar above denotes the total number of genes explained by that combination. The bars on the right indicate the total number of explained genes explained by each individual analysis (these are thus redundant with the columns representing a single dot). Notably, KLF4 ChIP-seq analysis (red box, and top-most bar on the right) alone already explained the majority of all explained genes.

Consistent with the increase in chromatin accessibility at CTCF TFBS in *Ago2&1*_KO cells (Fig 3D), we observed that the number of ChIP-seq-indicated CTCF-bound promoters and enhancers was higher for the upregulated *Ago2&1*_KO specific genes (35%, 260) than for the downregulated ones (19%, 201) (Fig 4B, Dataset EV5). While the ChIP-seq-based analysis for CTCF substantially increased the number of explained genes from 8% to 25% (compared to the motif-/ATAC-seq-based approach), interestingly, we found that a much larger fraction of the promoter and enhancer regions of *Ago2&1*_KO specific DEGs were bound by KLF4 (upregulated genes, 497, 73%; downregulated genes, 47%, 550) suggesting a major role in the regulation of *Ago2&1*_KO specific DEGs (Fig 4C, Dataset EV5).

In conclusion, the analysis of different genomics datasets allowed us to identify KLF4 as the major contributor to the *Ago2&1*_KO DEGs (Fig. 4D and E, Dataset EV5).

## DISCUSSION

In this work, we showed that AGO2&1 regulate specific gene expression in mESCs. The transcriptome analyses of *Ago2&1*_KO mESCs and other *miRNA* mutant mESCs indicated that the change in gene expression upon loss of AGO2&1 can be attributed only to a small degree by the loss of miRNA-mediated regulation, underlying a non-canonical function of AGO2&1 in mESCs. Here, we have studied potential mechanisms of how AGO2&1 can affect gene expression and identified KLF4 as the major mediator of AGO2&1 regulated genes.

Given that the miRNA effector proteins (DGCR8, DROSHA, DICER and AGO2&1), are involved in the canonical miRNA biogenesis pathway, one might assume that their knockouts would lead to similar transcriptomic perturbations. As we observed previously (Schäfer et al., 2021), DEGs in *Ago2&1*_KO showed strong similarities to *Dicer*_KO, but not to *Dgcr8*_KO or *Drosha*_KO. This might be partially attributable to noncanonical miRNA pathways, which function independently of the Microprocessor (DGCR8/DROSHA), but still require DICER and the AGO proteins (Bodak et al., 2017a). Thus, a limited set of miRNAs might still be active in *Dgcr8*_KO, *Drosha*_KO but not in *Dicer*_KO and *Ago2&1*_KO. Furthermore, AGO2 protein levels are strongly reduced in *miRNA*_KO mESCs due to its targeted proteasome degradation in the absence of miRNAs (Bodak et al., 2017b; Smibert et al., 2013). Some miRNA-independent AGO-mediated functions might therefore also be partially affected in *miRNA*_KO mESC lines. Surprisingly, despite these similarities, comparing DEGs between *Ago2&1*_KO and other *miRNA*_KO mESCs revealed a large number of DEGs that were specific to individual mutants and especially to *Ago2&1*_KO (Fig 1A), indicating potential miRNA-independent AGO specific functions. Indeed AGO specific functions, for example nuclear functions, have already been described in cancer cell lines and more recently also in mESCs (Li et al., 2020; Meister, 2013; Müller et al., 2021; Sarshad et al., 2018). Interestingly, we recently reported a global redistribution of H3K9me3 upon *Ago1* depletion in mESCs, however, this had very little impact on gene expression (Fig EV1A) (Müller et al., 2021). Similarly, *Ago2*_KO transcriptome also showed largely unchanged gene expression levels (Fig EV1A) (Ngondo et al., 2018). Given the strong observed perturbation in the *Ago2&1*_KO transcriptome, this might indicate that AGO1 and AGO2 do have global compensatory mechanisms in mESCs such that only disrupting the function of both proteins impacts gene expression strongly.

One such function is the regulation of SUZ12 and EZH2, two key components of PRC2. We showed that SUZ12 and EZH2 in *Ago2&1*_KO mESCs are reduced at protein levels but not at mRNA levels. As consequence, the amount of H3K27me3 in cells lacking AGO2&1 is globally reduced. However, the loss of H3K27me3 is not sufficient to fully explain the gene expression changes observed in *Ago2&1*_KO mESCs.

The integrative analyses of differential chromatin accessibility at predicted TFBS revealed a strong regulatory potential of two TFs, CTCF and, in particularly, KLF4, that can display both an activating and a repressive function (Bialkowska et al., 2017). Indeed, we found that KLF4 binds the promoter of about 50-70% of *Ago2&1*_KO specific up- and downregulated genes, respectively. Importantly, we also observed that the expression of KLF4 is strongly reduced in *Ago2&1*_KO mESCs. KLF4 is a pluripotency factor, which has been reported to occupy promoters of other key pluripotent transcription factors OCT4/SOX2/NANOG (OSN), thereby affecting their expression and *vice versa* (Bialkowska et al., 2017; Jiang et al., 2008). In addition, high KLF4 levels are associated with pluripotency and low levels with the onset of differentiation (Bialkowska et al., 2017). Accordingly, recent RNA-seq and Ribo-seq data (Schäfer et al., 2021) indicated that the pluripotency factors *Nanog* and *Oct4* are upregulated in *Ago2&1*_KO mESCs. Further, it has also been reported that in the absence of KLF4, the expression of two other Krüppel-life factors (KLF2, KLF5) increase, probably in order to compensate for KLF4 loss (Bialkowska et al., 2017; Di Giammartino et al., 2019). Indeed, we also observed an increase in the expression of *Klf2*, but not *Klf5*, in *Ago2&1*_KO mESCs (Dataset EV1). However, functional studies would be needed to confirm whether this increase potentially compensates for some of the loss of KLF4 in *Ago2&1*_KO. Thus, it might be interesting in the future to assess in detail the pluripotency status of these cells.

Interestingly, KLF4 has also been linked to other functions apart from regulating the pluripotency network. Recently, KLF4 has been shown to interact with SUZ12, a PRC2 complex member in mESCs (Di Giammartino et al., 2019). The observed reduction of SUZ12 at protein levels in *Ago2&1*_KO mESCs (Fig EV2A) suggests that the loss of KLF4 might destabilize SUZ12. This might be worth further investigations in the future, also with regard to the deposition of H3K27me3 in *Ago2&1*_KO mESCs, which then might be attributed to a secondary impact due to the loss of KLF4 in these mutant cells.

KLF4 has also been implicated in genome reorganization in mESCs and important functions in conferring enhancer connectivity (Di Giammartino et al., 2019). There, the disruption of KLF4 and its binding sites in mESCs showed an abrogation of enhancer contacts, which consequently decreased expression levels of associated genes (Di Giammartino et al., 2019). The loss of KLF4, along with many AGO2&1-regulated genes in our study, potentially may lead to alterations in chromatin conformation, which might be linked directly or indirectly to the AGO proteins. Indeed, previous studies already pointed towards potential alterations of the chromatin conformation linked to the AGO proteins (Moshkovich et al., 2011; Shuaib et al., 2019). In Drosophila, AGO2 has already been shown to interact with the architectural protein CTCF and to localize to chromatin regions, while abrogating this interaction led to reduced chromatin looping (Moshkovich et al., 2011). Also, in human cells, AGO1 was reported to be required for 3D chromatin maintenance. One study described that the loss of AGO1 leads to a disorganization of chromatin structure, which subsequently perturbed gene expression (Shuaib et al., 2019). Taken together, our observations as well as previous studies suggest a potential link for the AGO proteins in the modulation of chromatin conformation. Thus, mESCs might provide a unique opportunity to further investigate chromatin-related functions of the AGOs in mammals.

In conclusion, our study revealed non-canonical functions of AGO2&1 in mESCs that do not overlap with the canonical miRNA pathways and revealed a novel axis of gene regulation through the transcription factor KLF4.

## MATERIALS AND METHODS

### Mouse ESC lines

WT and all mutant E14 mESC lines (129/Ola background) were cultured as described in (Müller et al., 2021; Ngondo et al., 2018; Schäfer et al., 2021).

### CRISPR/Cas9 mediated gene knockout

The generation of the *Dgcr8_KO* mESCs and *Drosha_KO* mESCs cell lines were previously published by (Cirera-Salinas et al., 2017). The *Dicer_KO* mESCs lines were previously published by (Bodak et al., 2017b). The *Ago2&1_KO* mESCs were previously published by (Schäfer et al., 2021).

### Protein extraction and Western Blot Analysis

Protein extraction was performed in RIPA lysis buffer (50 mM Tris-HCL pH 8.0, 150 mM NaCl, 1% IGEPAL CA-630, 0.5% sodium deoxycholate, 0.5% sodium dodecyl sulfate supplemented with EDTA-free protease inhibitor cocktail (Roche)). The cell pellet was resuspended in ice-cold RIPA buffer and sonicated twice for 10 seconds. Afterwards samples were centrifuged 10 min at 10000 rpm and the supernatant was retrieved in a new 1.5 ml Eppendorf tube. The concentration of the protein extraction was assessed with a Bradford assay (Bio-Rad Laboratories). Laemmli buffer to a concentration of 1x was added to the samples and the samples were denatured for 5 min at 95°C. Western blotting has been performed as described in (Müller et al., 2021). Antibodies used for the WBs were: TUBULIN (Sigma-Aldrich T6199, 1:10000), H3K27me3 (ab6002, 1:2000), H3K27ac (ab177178, 1:2000), H3K4me3 (ab8580, 1:2000), H3K9me3 (ab8898, 1:2000), H3 (ab1791, 1:2000), SUZ12 (CST #3737), EZH2 (CST #5246), CTCF (ActiveMotif 61311, 1:2000), KLF4 (R&D AF3158, 1:2000).

### RNA extraction and quantitative RT PCR Analysis

The RNA extraction has been carried out as described in (Bodak and Ciaudo, 2016). Briefly, RNA extraction from cell pellets was performed with the Trizol reagent (Life Technologies). 1ml Trizol was added to the cell pellet and mixed by vortexing for 10 seconds. Then, 200 μl Chloroform (Sigma-Aldrich) was added, and samples were vortexed again for 10 seconds. Samples were centrifuged at 4°C for 15 min at 13500xg and the upper phase was transferred into a new 1.5 ml Eppendorf tube. 600 μl isopropanol (Merck) was added, mixed by vortexing and samples were centrifuged for 30 min at 4°C at 13500xg. The supernatant was removed, and the RNA pellet was washed once with 1 ml ice-cold 70% Ethanol and centrifuged again for 10 min at 13500xg. The supernatant was removed, the RNA pellet was air-dried and resuspended in RNase-free water. The quality of 1 μg of RNA was checked by on a 1% agarose gel (Sigma). RT and qPCR has been performed as described by (Müller et al., 2021). Primers are listed in Table EV6.

### Reference genome and gene annotation

All OMICs analyses were based on the mouse reference genome *GRCm38/mm10*. ChIP-seq analyses were performed using GENCODE mouse gene annotations in version 23 (Frankish et al., 2019) and Drosophila genome in version 6 (Hoskins et al., 2015) for spike-in normalization, while all other analyses were performed using the comprehensive mouse gene annotation file from ENSEMBL in version 98 (Cunningham et al., 2019).

### RNA-seq analysis of RNAi mutants

RNA-seq data was obtained from GEO (Table EV6) and analyzed as described in (Schäfer et al., 2021). Briefly, the snakePipes *RNA-seq* pipeline (Bhardwaj et al., 2019) was employed with default arguments for trimming, quality control, mapping, read counting and differential expression analysis using *Trim Galore/cutadapt* (Martin, 2011), *FastQC/multiQC* (Andrews et al., 2012), *Bowtie2* (Langmead and Salzberg, 2012), *featureCounts* (Liao et al., 2014) and *DESeq2* (Love et al., 2014) respectively.

#### Definition of Ago2&1_KO-specific DEGs

DEGs in each of the *miRNA*_KO mutants were defined with an adjusted p-value threshold of 0.05 and a minimal log2FoldChange of 0.5. The set of *Ago2&1*_KO-specific misregulated genes described throughout this paper (Dataset EV1) consists of genes that are differentially expressed in *Ago2&1*_KO and *not* differentially expressed in any of the other *miRNA*_KO mutants (*Dgcr8*_KO, *Drosha*_KO, *Dicer*_KO). Visualizations of differential expression and gene set overlaps were realized as MA-plots and Venn-diagrams using matplotlib/seaborn (Hunter, 2007; Waskom, 2021) and eulerr/matplotlib_venn (Larsson et al., 2018, http://eulerr.co/) respectively.

For the gene ontology analysis, the ClueGO App (Bindea et al., 2009) for Cytoscape (Shannon et al., 2003) was used. For the analysis, the terms ‘Biological Process’, ‘Molecular Function’ and ‘Cellular Component’ were used (date: 2020/03/09). Only pathways with pValues < 0.005 were considered and Terms/Pathways with a minimum GO tree interval level of 3 and a maximum of 10 were used. To represent the size of the nodes according to their pValues, the Bonferroni step down corrected Term PValue was selected and the −log10 of these pValues was calculated and represented in the figure.

### Analysis of histone mark levels for gene groups

Gene groups were defined as follows. *Ago2&1*_KO specific DEGs were split based on their differential expression into an *upregulated* and a *downregulated* group. *Predicted MiRNA Targets* were taken as provided by (Schäfer et al., 2021). Expressed genes were defined as genes from the comprehensive gene annotation set from ENSEMBL (described above) with a minimum expression of 1 TPM in our WT RNA-seq data (Dataset EV1). For each gene, all annotated transcripts were considered as separated instances. Each transcript was converted to a range of 10 kbp around the transcription start site in order to account for the diverse placement patterns of different histone marks.

Genome-wide histone mark levels for seven different marks were obtained from ENCODE (experiment IDs ENCSR000CGO, ENCSR000CGN, ENCSR000CGQ, ENCSR000CGP, ENCSR000CGR, ENCSR000ADM, ENCSR059MBO, Davis et al., 2018; Dunham et al., 2012) as bigwig files of signal over input log2FoldChanges for two combined replicates. For each region in each of the described groups, the mean signal was obtained for each histone mark using the *multiBigwigSummary* tool (Ramírez et al., 2016). Finally, the mean signals were averaged across all transcripts of each gene group and z-score normalization was applied on a per histone mark basis.

### H3K27me3 ChIP-seq

#### Sample and library preparation

The chromatin extraction and pull-down has been performed as described in (Müller et al., 2021). In order to prepare chromatin for sequencing, two replicates have been performed, where 100 μg of chromatin, together 125 ng Drosophila spike-in chromatin (a kind gift from the Santoro lab), was precleared with 50 μl of Dynabeads protein G (ThermoFisher Scientific) and further used for the chromatin pull-down. 10 μg of H3K27me3 (ab6002) was used for the pull-down.

Library preparation and sequencing was performed by the Functional Genomics Center Zürich (FGCZ). For the library preparation the NEBNext® Ultra™ II DNA Library Prep Kit from NEB (New England Biolabs, Ipswich, MA) was used. 1 ng starting material of each sample was used, end-repaired and afterwards adenylated. Then, indexed adapters were ligated to the fragmented samples and PCR was performed to enrich the fragments. The library quality and quantity were assessed by with the help of a Qubit® (1.0) Fluorometer and a Tapestation (Agilent, Waldbronn, Germany). The libraries were sequenced on the Illumina Novaseq 6000 (Illumina, Inc, California, USA) by 100 bp single reads.

#### ChIP-seq analysis pipeline

Adaptors were trimmed using *trimmomatic* (Bolger et al., 2014). Sequencing reads were mapped to the mouse mm10 genome and to the Drosophila dm6 genome using *bowtie2* (Langmead and Salzberg, 2012). Duplicates were marked and filtered out using *samtools* (Li et al., 2009). *bamCompare* from deepTools (Ramírez et al., 2016) was used to generate a bigwig file of IP over Input. Drosophila-mapped read counts were used as scaling factor for the Input and the IP samples in order to get normalized read counts.

#### Cluster-analysis of transcript-centric histone modification levels

Both WT ChIP-seq replicates were compared to the WT H3K27me3 ChIP-seq data from ENCODE (ENCSR059MBO, Davis et al., 2018; Dunham et al., 2012). Compared to the already published ENCODE ChIP-seq, one replicate exhibited stronger signal-to-noise ratio and this sample and the according *Ago2&1*_KO replicate were selected for representative images and cluster analysis in Fig. 2 and EV2.

The published H3K27me3 ChIP-seq data from ENCODE (ENCSR059MBO, as described above) was compared to our generated data for experimental validation. To improve comparability and account for differences in samples preparation and data analysis between the published and our data, signal levels of the ENCODE data were normalized to our WT sample (scaling with a factor of 1/3). Clustered heatmaps with mean profiles for four clusters were generated using deeptools’ *computeMatrix* and *plotHeatmap* tools for the complete set of annotated transcripts around TSS (<5 kbp distance). Transcripts from each cluster were reduced to their corresponding genes and associated with differential expression in *Ago2&1*_KO for comparison of log2FoldChange distributions across the four clusters using CDF curves plotted with matplotlib/seaborn (Hunter, 2007; Waskom, 2021).

For the comparison of the H3K27me3 ChIP-seq cluster_1 with known bivalent genes, the list of high-confidence bivalent “HC-Bivalent” genes from (Asenjo et al., 2020) was retrieved.

The *pyGenomeTracks* tool was used in order to show a genome browser view of selected regions (Ramírez et al., 2018).

### ATAC-seq

#### Sample & library preparation and sequencing

Two millions of cells were trypsinized and resuspended in 1 ml of freezing medium (serum + 10% DMSO). Cells were transferred into a freezing container and stored at −80°C. ATAC-seq sample preparation, library preparation and sequencing have been performed by Quick Biology (https://www.quickbiology.com/ngs-services/ATAC-seq-service) according to (Corces et al., 2017) with 50,000 intact cells. Cells were first washed, then lysed to obtain nuclei preparation. Using aTn5 transposase, the genome was simultaneously fragmented and tagmented, leading to amplifiable DNA fragments with sequencing adapters for the Illumina platform. The fragments were amplified by PCR and purified using Qiagen MinElute PCR Purification Kit (Qiagen, Maryland, USA). The final library quality was controlled using the Agilent Bioanalyzer 2100 (Agilent Technologies, Santa Clara, CA) and the quantity assessed using the Life Technologies Qubit 3.0 Fluorometer (Life Technologies, Carlsbad, CA). Finally, the libraries were sequenced on an Illumina HiSeq X Ten Sequencer (Illumnia Inc., San Diego, CA) with 2 x 150 bp read (paired end).

#### ATAC-seq analysis pipeline

Raw read counts were processed by the snakePipes *DNA-mapping* and *ATAC-seq* pipeline (Bhardwaj et al., 2019). Briefly, reads were trimmed using *Trim Galore/cutadapt* (Martin, 2011) with parameters “*–nextera –paired*” after which *Bowtie2* was called for mapping with default arguments. Reads were filtered for PCR duplicates and mappings were filtered if mapping quality was below “3” or if the fragment size was smaller than 150 bp, to only keep reads originating from nucleosome free regions (NFRs). *Genrich* (available at https://github.com/jsh58/Genrich) was used for peak calling and CSAW/EdgeR (Lun and Smyth, 2016) determined Differential Accessibility (DA) of the called peak regions. Analysis of quality control metrics indicated differences in experimental ATAC efficiencies across samples. Therefore, fraction of reads in peaks (FRiPs) score was used as linear scaling factors in downstream analyses (differential accessibility, MA plots, heatmap/profile plots, diffTF analysis (see below)) to compensate for differences across samples, as suggested by (Reske et al., 2020).

Comparison of WT and *Ago2&1*_KO were performed using (i) deeptools heatmap/profile plots over gene regions, similarly as described in the H3K27me3 ChIP-seq analysis, (ii) profile plots at promoter regions (<3 kbp TSS distance of all annotated transcripts) using deeptools *computeMatrix* and *plotProfile* (iii) profile plots at enhancer regions, as determined by (González-Ramírez et al., 2021) (iv) MA plots of the differential accessibility at peak regions (using matplotlib/seaborn, Hunter, 2007; Waskom, 2021).

#### Integration with RNA-seq and TF motifs

Peaks with statisticaly significant DA (p<0.05) were associated with genes that were in 1000 bp proximity to their transcription start sites. Genes associated with increased chromatin accessibility and decreased accessibility were grouped separately and associated with differential expression as assessed by RNA-seq, which allowed for a comparison of their log2FoldChange distributions (CDF plots were generated using matplotlib/seaborn, Hunter, 2007; Waskom, 2021),

The *diffTF* tool (Berest et al., 2019) was employed to assess potential differential binding of Transcription Factors (TFs), based on chromatin accessibility data. The tool relies on previously determined binding motifs for a large set of transcription factors and assesses DA at predicted motif-based TF binding sites (TFBS) within the genome. Per-TFBS information and per-TF summary data is generated and indicates whether TF binding sites were associated with increased or decreased DA. diffTF was run on all TFs having minimal expression of 1 TPM in our WT RNA-seq data and with a binding motif available as obtained from the HOCOMOCO database (v10, Kulakovskiy et al., 2018).

TFBS-level information was obtained for the 5 TFs with most significant DA at TFBSs, filtered for statistical significant DA (p < 0.05) and associated with genes in two ways: (i) promoter-based association was performed using *bedtools closest* (Quinlan and Hall, 2010) on gene TSSs with a 1000 bp distance filter, (ii) enhancer-based association was performed based on enhancer-gene associations as provided by (González-Ramírez et al., 2021, “Hi-C–top” active and poised enhancers were combined); TFBSs that were overlapping with reported enhancer regions were associated with the corresponding genes. Associated DA and DEG information was then visualized on a per-gene basis using scatterplots (Hunter, 2007; Waskom, 2021).

### Integration of published KLF4- and CTCF-ChIP-seq data

CTCF ChIP-seq peaks were obtained from (Nora et al., 2017). KLF4 ChIP-seq peaks were obtained from (Di Giammartino et al., 2019), where all reported peaks but “Transient”-labeled ones were considered.

Peaks were associated with genes again by promoter proximity (<1000 bp to TSS) and enhancer overlap (González-Ramírez et al., 2021) similarly to as indicated in the last section, but using pybedtools (Dale et al., 2011; Quinlan and Hall, 2010). Associated genes were filtered for DEGs in *Ago2&1*_KO and used for Venn diagram overlaps with previously defined gene groups.

### Combined contribution of regulatory mechanisms to Ago2&1_KO-specific DEGs

Gene sets derived from the previously described mechanisms (H3K27me3, chromatin accessibility, general TF-binding, CTCF- and KLF4-binding) were intersected with the set of *Ago2&1*_KO-specific DEGs and overlapped using an UpSet plot in a union-fashion, as implemented by (Gu et al., 2016).

## QUANTIFICATION AND STATISTICAL ANALYSIS

See Methods Details on how quantification and statistical analyses have been performed. If not mentioned otherwise, statistical analysis has been performed using PRIMS 8 more detail is indicated in the figure legends.

## DATA AVAILABILITY

Data visualization has been performed with the tools, described in the Method sections (Table EV6). If not mentioned otherwise graphs have been generated by using PRIMS.

All code for data analyses and visualizations described in the paper are found in the following github repository https://github.com/moritzschaefer/ago21_specific_effects.

Sequencing data has been deposited on GEO (Table EV6)

The ATAC-seq and ChIP-seq data has been deposited to the GEO database: GSE185410 Reviewer access token: gnszaiwsbrunpwt

## Acknowledgments

We would like to thank the members of the Ciaudo lab and Dr. Tobias Beyer for fruitful discussions and the critical reading of this manuscript. This work was supported by the Swiss National Science Foundation (grants 31003A_173120 and 310030_196861) to C.C. In addition, C.C, M.M and R.S were supported by the NCCR RNA and Disease. We also want to thank the Functional Genomics Center Zurich (FGCZ) for their support with the preparation of ChiP-seq libraries and sequencing. We also want to thank Quick Biology for their help with the ATAC-seq library preparation, sequencing and computational support.

## Author Contributions

Conceptualization, MM, MS and CC; laboratory experiments, MM, TF; ChiP-seq libraries preparation, MM, RPH; computational analysis, MS and DS; writing original draft preparation, MS, MM and CC; writing, review and editing, CC; expertise and editing, RS; visualization, MS, MM and CC.; supervision, CC; funding acquisition, CC. All authors have read and agreed to the published version of the manuscript.

## Conflict of Interests

The authors declare no financial and non-financial competing interests.

## Supporting Information

Expanded View Figures

Dataset EV1

Dataset EV2

Dataset EV3

Dataset EV4

Dataset EV5

Table EV6

---

**Figure EV1.**
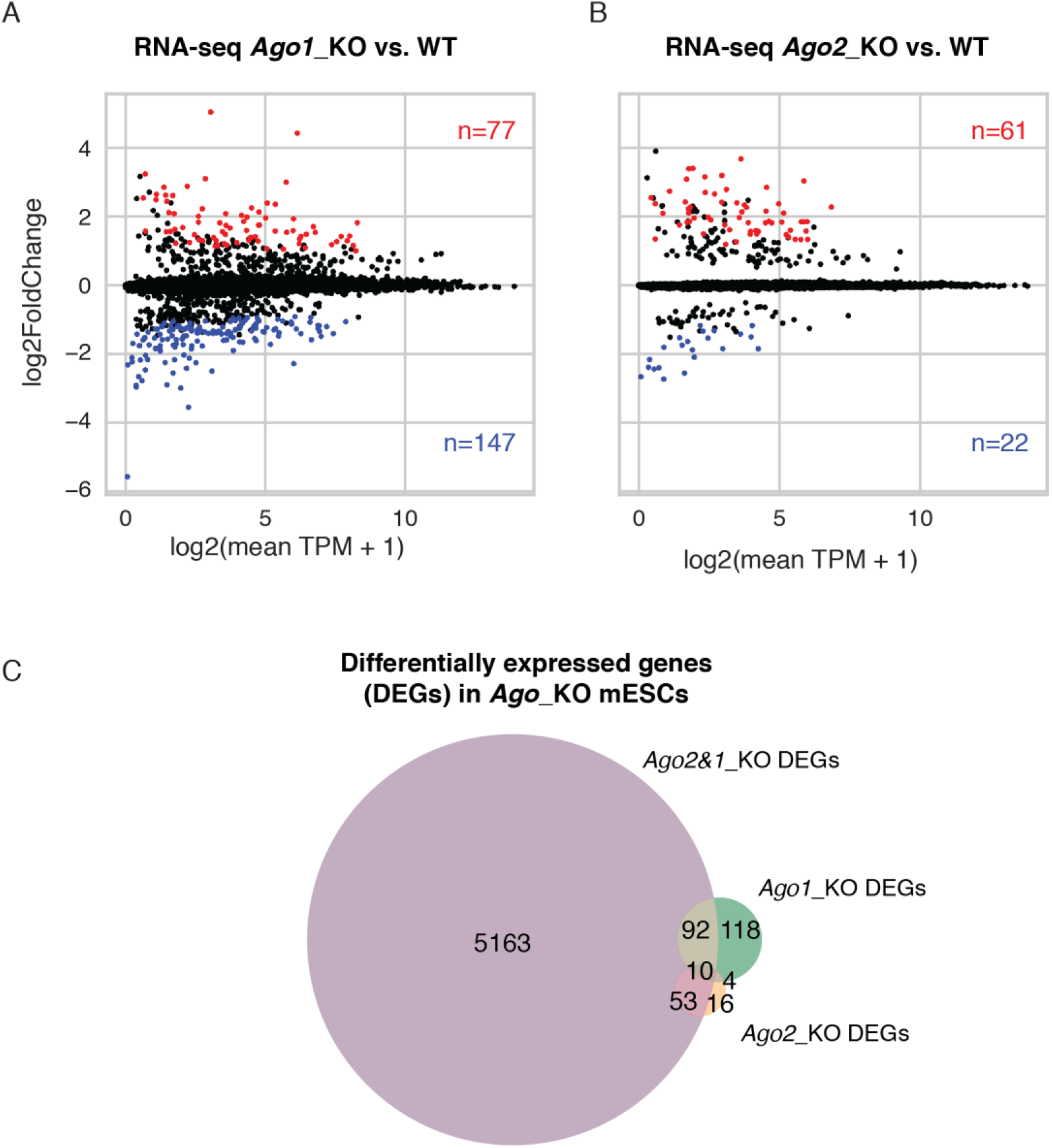
*Ago*_KO transcriptomic analysis. (A, B) MA plots showing the differential gene expression for the *Ago1*_KO mESCs versus WT as in (Müller et al., 2021) (A) and *Ago2*_KO vs WT as in (Ngondo et al., 2018) (B). Highlighted in red are the statistically significant upregulated and in blue the statistically significant downregulated genes. The number of genes in those groups are denoted in the right corners (Dataset EV1). TPM: transcripts per Million. (C) Venn diagram showing the overlap between differentially expressed genes (DEGs) in *Ago2&1*_KO, *Ago1*_KO and *Ago2*_KO mESCs. Notably, *Ago1*_KO and *Ago2*_KO show comparably low DEGs and overlap only partially with *Ago2&1*_KO.

**Figure EV2.**
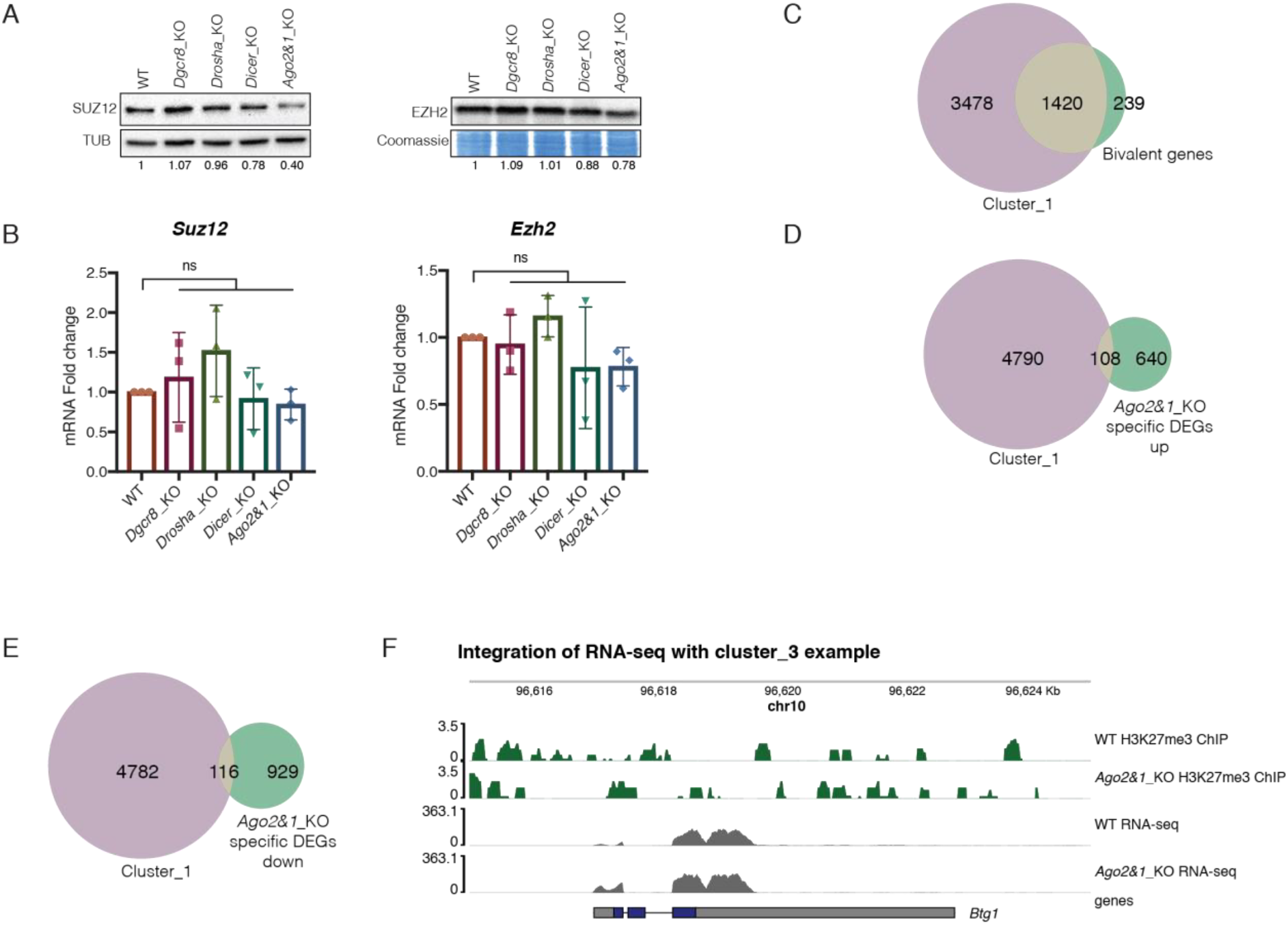
Expression of *Suz12* and *Ezh2* PRC2 members in *Ago2&1*_KO mESCs and the integration of the *Ago2&1*_KO transcriptome with the H3K27me3 ChIP-seq. (A) Representative Western blots for SUZ12 and EZH2 in WT, *Dgcr8*_KO, *Drosha*_KO, *Dicer*_KO and *Ago2&1*_KO mESCs out of n=3 independent experiments. Tubulin (TUB) and Coomassie were used as a loading control. Quantification for each individual experiment is shown below the blot. (B) Quantitative-RT-PCR results for *Suz12* and *Ezh2* in WT, *Dgcr8*_KO, *Drosha*_KO, *Dicer*_KO and *Ago2&1*_KO, from n=3 independent experiments. ns = non-significant, unpaired t-test. (C) Venn diagram showing the overlap of genes in the ChIP-seq cluster_1 with high-confidence bivalent genes, identified by (Asenjo et al., 2020). (D, E) Venn diagrams showing the overlap of genes in the ChIP-seq cluster_1 with *Ago2&1*_KO specific up (D) and down (E) DEGs. (F) Genome browser view of an example gene from the ChIP-seq cluster_3 (Btg1), showing both ChIP-seq and RNA-seq coverages. Depicted are genome tracks for the WT and *Ago2&1*_KO ChIP-seq (one sample per experiment and condition). The y-axis represents the deposition of H3K27me3 (for the ChIP-seq in green) or the expression levels of the RNA-seq in gray.

**Figure EV3.**
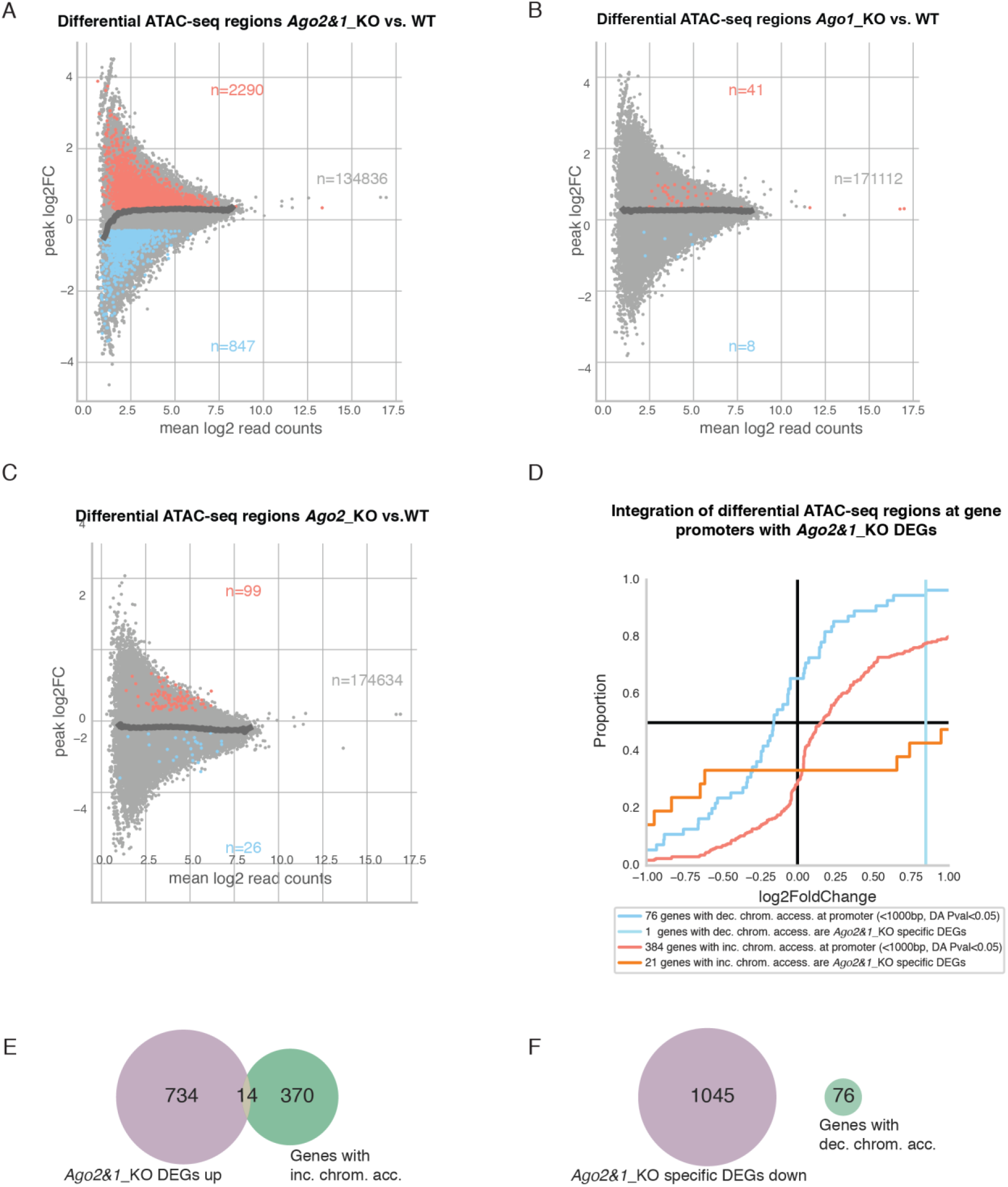
Integration of the *Agos*_KO transcriptome with chromatin accessibility. (A, B, C) MA plots of genomic regions with detectable chromatin accessibility (based on Genrich peak calling). Differential accessibility (DA) at those regions was assessed using CSAW/EdgeR for *Ago2&1*_KO vs WT (A), *Ago1*_KO vs WT (B) *Ago2*_KO vs WT (C). Regions with an absolute log2FC > 0.3 and adjusted p-value < 0.05 were considered as statistically significant and are colored (red and blue) and counted. (D) Cumulative Distribution Function (CDF) plot of differential expression (DE, log2FoldChange of RNA-seq *Ago2&1*_KO vs WT) for genes with promoters in proximity (<1000bp) of differentially accessible regions (p-value <0.05) with increased (blue) or decreased (red) accessibility. DEGs were additionally shown after filtering by *Ago2&1*_KO specific DEGs. (E, F) Overlap of genes associated with differential accessibility (as in (D)) with *Ago2&1*_KO specific up (E) and down (F) DEGs.

**Figure EV4.**
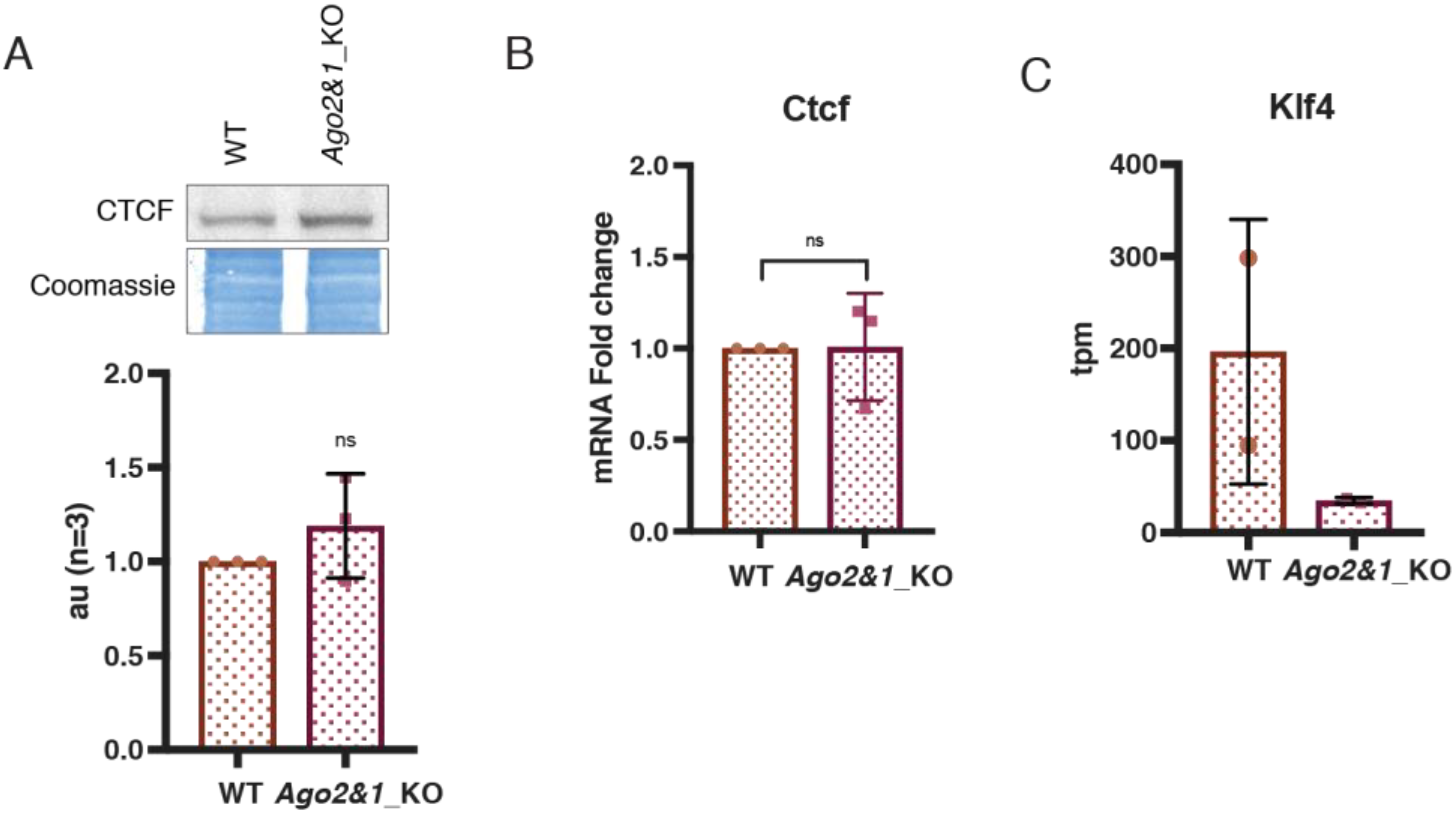
Characterization of CTCF and KLF4 in mESCs. (A) Representative Western blot (top) for CTCF in WT and *Ago2&1*_KO mESCs and quantification (bottom) out of n=3 independent experiments, ns=not significant, unpaired t-test. (B) *Ctcf* qRT-PCR results in WT and *Ago2&1*_KO mESCs. n=3 independent experiments. ns=not significant, unpaired t-test. (C) *Klf4* tpm counts from the RNA-seq data (Dataset EV1) in WT and *Ago2&1*_KO mESCs.

